# Explaining Missing Heritability Using Gaussian Process Regression

**DOI:** 10.1101/040576

**Authors:** Kevin Sharp, Wim Wiegerinck, Alejandro Arias-Vasquez, Barbara Franke, Jonathan Marchini, Cornelis A. Albers, Hilbert J. Kappen

## Abstract

For many traits and common human diseases, causal loci uncovered by genetic association studies account for little of the known heritable variation. Such ‘missing heritability’ may be due to the effect of non-additive interactions between multiple loci, but this has been little explored and difficult to test using existing parametric approaches. We propose a Bayesian non-parametric Gaussian Process Regression model, for identifying associated loci in the presence of interactions of arbitrary order. We analysed 46 quantitative yeast phenotypes and found that over 70% of the total known missing heritability could be explained using common genetic variants, many without significant marginal effects. Additional analysis of an immunological rat phenotype identified a three SNP interaction model providing a significantly better fit (p-value 9.0e^-11^) than the null model incorporating only the single marginally significant SNP. This new approach, called GPMM, represents a significant advance in approaches to understanding the missing heritability problem with potentially important implications for studies of complex, quantitative traits.

---

The problem of ‘missing’ (or hidden) heritability^1^ is a central challenge for the genetics of complex traits. In human populations, genome-wide association studies (GWAS) have successfully identified over 1,100 genetic loci associated with over 165 common traits, but the sum of their independent effects often appears to explain only a small proportion of the total heritable variation (broad-sense heritability)^2,3^. An important open question is how much additional heritability might be explained by the non-additive effects of interactions between genetic loci (epistasis). Arguments for the ubiquity and importance of epistasis^4,5,6^ are supported both by evidence from empirical studies in model organisms such as mouse^7^, yeast^8,9^ and drosophila^10^ and also by demonstrations that a broad class of interaction models are capable of explaining any amount of heritability while exhibiting minimal marginal effects for individual loci^11,12^. However, while progress has been made in estimating the phenotypic variation that might be explained by genetic variants in a purely additive model^13^(narrow-sense heritability), methods for estimating broad-sense heritability directly from common genetic variants are lagging behind.

Detecting non-additive, interaction effects of genetic variants is difficult for two principal reasons: possible models are of unknown order and complexity and the number of loci considered is typically high. Parametric approaches that test the likelihood of a set of specified models face an exponential scaling of computational cost with interaction order as well as the problem of correcting appropriately for the vastly increased number of hypothesis tests. Current approaches^14-21^ constrain either the space of models (for example by limiting the search to test only for pairwise effects) or the set of genetic loci by some criterion, such as filtering based on lower order effects. However, in the presence of higher order interactions, this strategy results in reduced power to identify important loci and, hence, to explain variance arising from non-additive effects (non-additive variance).

Here we employed Gaussian Process Regression (GPR)^22^ to address these issues. GPR places no constraints on the order of possible interactions and makes minimal assumptions about their possible form. It models a probability distribution over the unknown regression function mapping genotype markers to a quantitative phenotype as a stochastic process. The effect of each marker is controlled by a scale hyperparameter of the process: smaller values imply greater sensitivity of the phenotype to changes in allele dosage. Other hyperparameters determine the partitioning of phenotypic variance between the model and unexplained noise. Inference is carried out using an efficient form of Markov Chain Monte Carlo (MCMC) sampling algorithm termed Hybrid Monte Carlo (HMC)^23^ Posterior probability distributions over the values of the hyperparameters, obtained via HMC simulations, can be used to estimate quantities of interest (such as the fraction of variance explained and the strength of evidence for association of each marker) and to predict phenotypes for unseen genotypes (**Methods**).

A useful way to think about the approach is by comparing it to methods, such as GCTA^24^, which estimate the narrow-sense heritability under an additive model. In GCTA, a single fixed kinship matrix is used to model a random effect term in a mixed model. The estimate of the variance component of this random effect is used directly to estimate heritability. In contrast, our method *learns* the kinship matrix, where the contribution of each marker to the kinship matrix is controlled by a set of parameters and occurs in a non-linear way. We call our method Gaussian Process Mixed Model (GPMM).

Accurate estimation of an arbitrary function of even a modest number of markers requires, in general, an impractically large number of samples. Nevertheless, we show that our approach has power to estimate non-additive variance and to identify important loci with no marginal effects even when there is insufficient data to estimate such a function for all possible genotypes. This power derives from a combination of three factors. Two are inherent properties of GPR: averaging over the uncertainty in plausible regression functions, and using a sparsity-inducing prior effectively embodying a prior skepticism about the importance of any specific locus (**Methods**). The third factor is the availability of biological replicates. Importantly, we found these significantly improved the power of GPR. In contrast, they made little difference to the results of linear modeling.

## RESULTS

### Accounting for non-additive variance

To demonstrate the utility of GPR, we firstly applied the method to published data on growth rates of 1,008 closely related strains of yeast under 46 different growth conditions^8^. REML estimates of broad-sense heritability (*H*^2^) using replicated segregants (**Methods**) for a number of these traits were significantly greater than those for narrow-sense heritability (*h*^2^). Furthermore, very little of this missing heritability could be explained by pairwise interactions^8^. Highly correlated single nucleotide polymorphisms (SNPs) were removed to minimize the number of essentially redundant explanations of the data (**Methods**), and GPR estimates of the broad-sense heritability 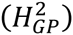 attributable to SNPs were calculated from the output of the MCMC sampler (**Methods**).

For 25 out of 46 phenotypes, where *H^2^* was high (≥ **70%**), **there was very good agreement with** 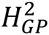 (**Fig. 1a**); the explanatory models found by GPR using SNPs explained all of the missing heritability. Both *H*^2^ and 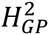 were higher than *h*^2^.

**Figure 1.**
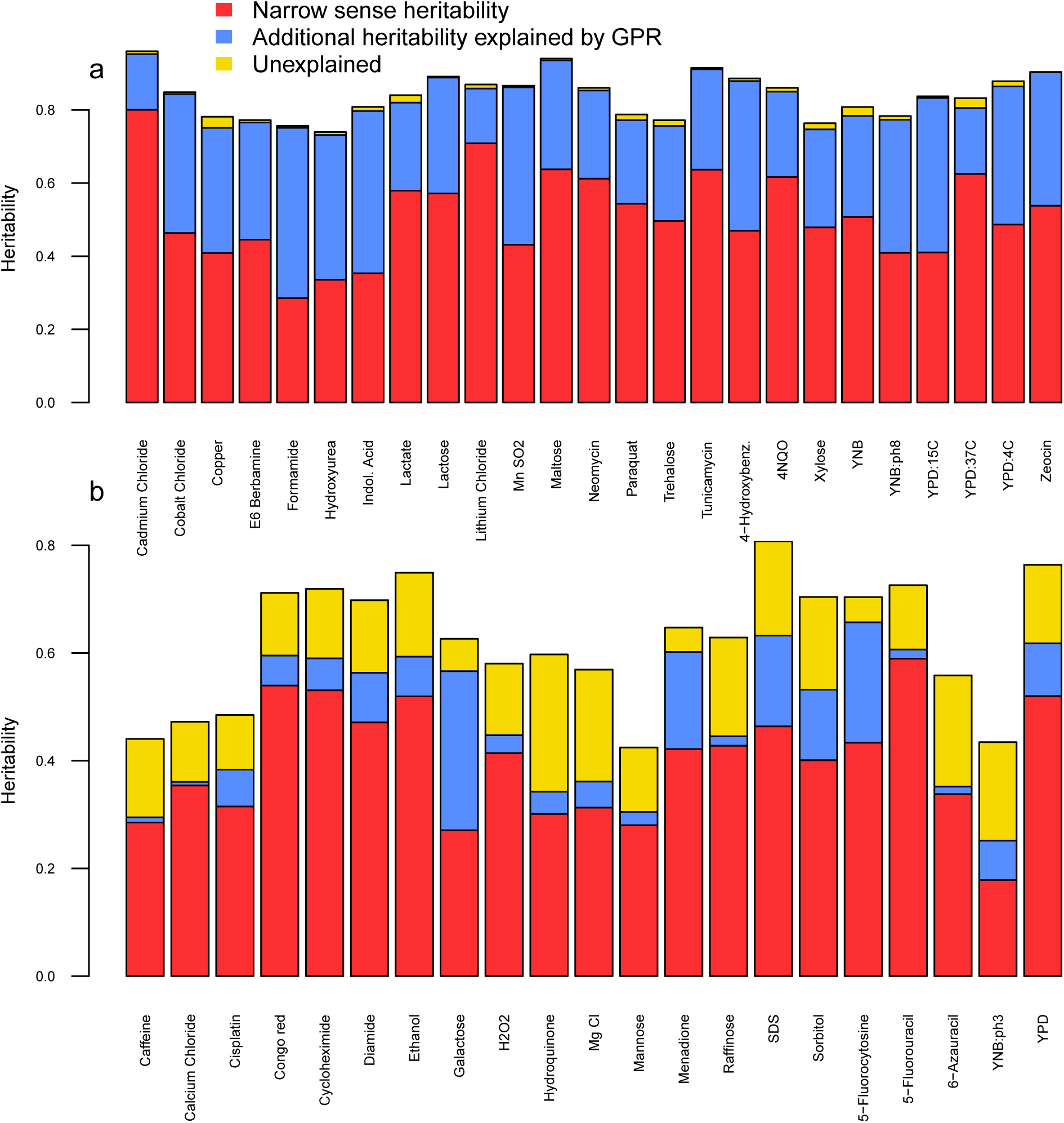
Comparison of heritability estimates. Each bar represents one of the 46 yeast growth conditions‥ The total height of a bar represents the broad sense heritability (*H*^2^); the heights of the different coloured subdvisions indicate the partioning of variance between three components: red bars correspond to the proportion of variance that is additive (*h*^2^); blue bars indicate the additional proportion of variance explained by GPR, i.e., *H*^2^_*GP*_ - *h*^2^; yellow bars correspond to residual variance. **a** The 25 growth conditions with estimated *H*^2^ ≥ 70% **b** The remaining 21 growth conditions. Estimates of *h*^2^ were obtained using only the pruned subset of markers used for inference by GPR. Supplementary Figure 2 shows the equivalent comparison for estimates of *h*^2^ obtained from the full set of 11,623 markers.

For the remaining 21 yeast growth conditions, 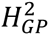 was apparently unable to explain missing heritability in its entirety. Nevertheless, even when 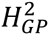 was lower than *H*^2^, they were never lower than estimates of narrow-sense heritability *h*^2^ from the same pruned subset of SNPs (**Fig. 1b**).

The ability of GPR to explain missing heritability was correlated with *H*^2^ (Pearson correlation = 0.82). (**Supplementary File 2**, **Fig. 1**) This was consistent with the standard errors of 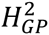, indicating that GPR inference of explained variance also becomes more uncertain as REML estimates decrease. As an illustration of this effect, there was clear evidence of two modes in the posterior distribution of explained variance (**Supplementary File 2**, **Fig. 3**) for the two conditions with the largest uncertainty (Ethanol and Hydroquinone). In one of these regions, GPR estimates were comparable to those of REML estimates of *H*^2^; in the other they were comparable to those of *h*^2^.

Comparison of 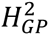 to reported estimates of *h*^2^ obtained from the full set of 11,623 markers^8^, gave similar results, but with a small number of exceptions. For four of the growth conditions (5-Fluorouracil, Hydrogen Peroxide, Congo Red and Cycloheximide) GPR heritability estimates were less than reported estimates of *h*^2^ (**Supplementary File 2**, **Fig. 1c & 2b**). We consider the interpretation of this observation below (**Discussion**).

### Gaussian Process Regression does not overfit

Given the flexibility of GPR, it is natural to ask whether it infers better models of the data or merely adapts to noise. We checked for overfitting in two different ways. Firstly, we examined the ability of GPR to predict phenotypes of held-out individuals. For each trait, we created random partitions of the data: 90% of the samples were used for training and the remaining 10% for testing. We found that the performance of GPR as assessed by the standardised mean squared error (SMSE) - the mean squared error normalised by the empirical variance of the data (**Methods**) – was never worse than that of linear regression (**Fig. 2**). This confirmed that GPR did not overfit.

**Figure 2.**
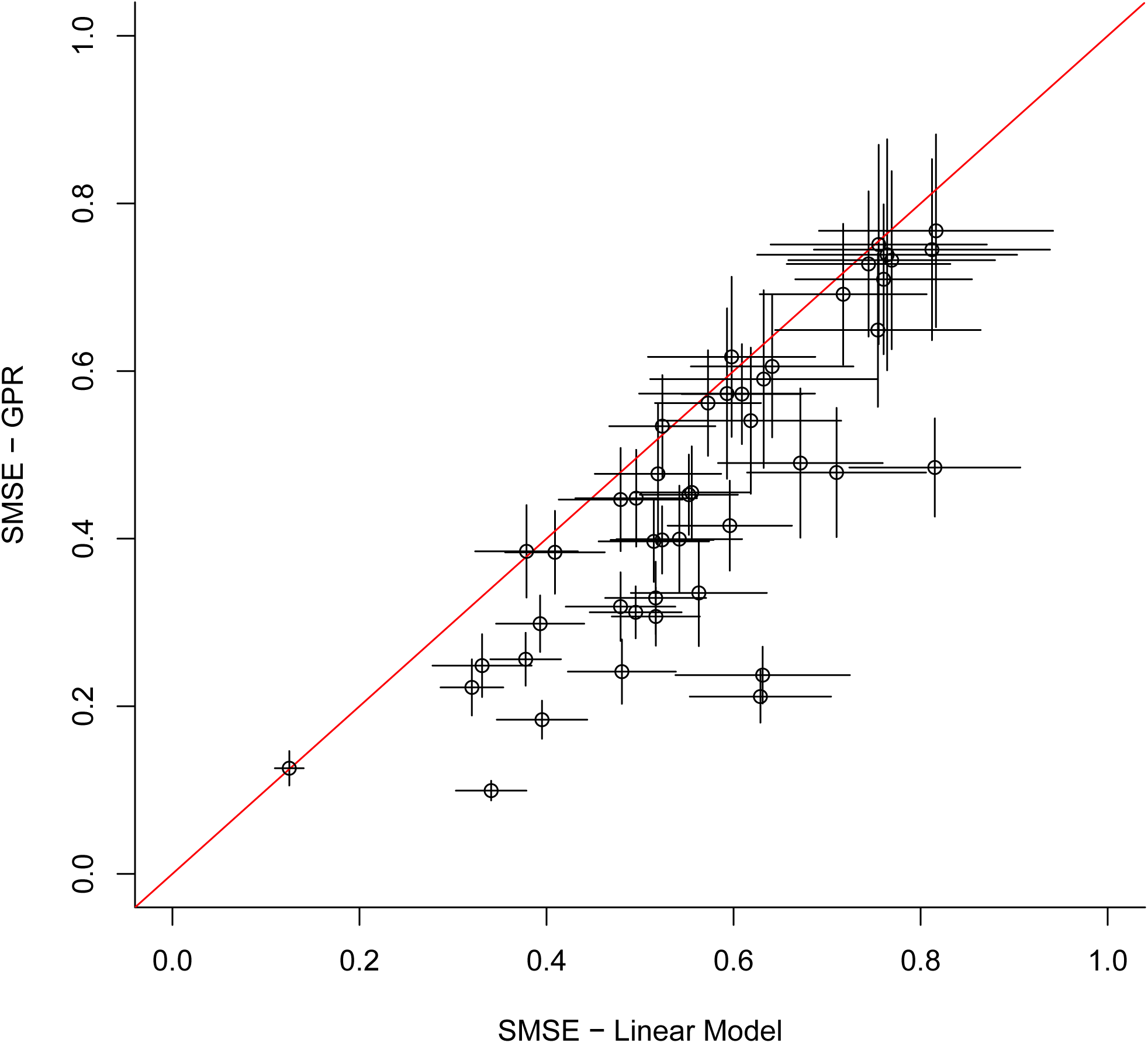
Comparison of standardised mean squared errors (SMSE) in predicted phenotypes made by GPR with those of a linear model obtained from multiple linear regression using the same SNPs. Each point corresponds to a held out sample on one of the 46 yeast growth conditions. Error bars correspond to ± 1 SE. The errors made by GPR predictions are systematically lower than those made by the linear model indicating that GPR is not overfitting. In this sense, GPR is finding better explanatory models than a purely additive model.

As an additional check, we applied both GPR and linear regression to simulated datasets consisting of real yeast genotypes combined with phenotypes generated from a purely additive model; the coefficients of this model were those obtained from the linear regression fit using the real phenotypes and only the additively significant subset of SNPs. We set *H*^2^= *h*^2^=0.5. GPR heritability estimates were consistent with both the underlying ground truth and those of the best fitting linear model while prediction errors for phenotypes of unseen test data were never worse than those of the linear model (**Supplementary File 2, Table 1**). Although able to create more complicated models, apparently explaining more heritability than a linear model, GPR does not do so when the true generative model is additive.

### Identifying new relevant loci

A key feature of our method is that we can infer the relative contribution of each locus to the model, in an analogous way to estimating the additive effect of a locus or its p-value of association. Relevance of a locus was determined from the marginal posterior distribution of the corresponding scale hyperparameter; a smaller scale implied greater relevance (**Methods**). These distributions were used to compute Bayes Factors (BF) to assess the strength of evidence for association of a given locus after averaging over possible models. (**Methods**). Values of 2, 6 and 10 for twice the natural logarithm of the Bayes Factor are sometimes taken to indicate positive, strong and very strong evidence respectively^25^.

For many conditions there was clear evidence that the sampler was exploring multiple modes of the posterior distribution corresponding to different explanations of the data. Different modes did not always incorporate the same subset of markers. Therefore, it was important to have confidence that the relative probability mass in the different modes had been correctly estimated. This was done by checking that different randomly initialized MCMC runs were converging to the same distribution using standard criteria (**Methods**). For only one condition (Manganese Sulphate) out of 46 was convergence not indicated in the time we allowed.

Illustrations of the power of GPR to detect higher order interactions are provided by yeast growth in the presence of Zeocin and Lactose. Missing heritability was estimated as ∼35% and ∼30% respectively, but there was no evidence for significant pairwise interactions in either case^8^. For Zeocin, GPR explained all of the missing heritability (**Fig.1** **and Supplementary File 1**), and found positive evidence for association (2log(BF) ≥2) for the 17 reported additive QTLs and a further 11 loci (**Fig. 3a** **and Supplementary File 1**). For two of these additional loci – Chr2: 700731 and Chr4: 832287 (loci 7 and 16 respectively in Figs. 3a and 3b) evidence of explanatory relevance was strong (2log(BF)≥6). For Chr4: 832287 in particular, the evidence (Pr(scale ≤ 10)=0.91) was stronger than that for many of the additive QTLs even though it ranked 57^th^ out of 59 in terms of effect size in a multiple linear regression model. These results cannot be due to LD with additive QTLs; no additive QTLs were identified on chromosome 4 and the pairwise correlation between Chr2: 700731 and the single additive QTL on the same chromosome, Chr2: 479195 was low (∼0.11).

**Figure 3.**
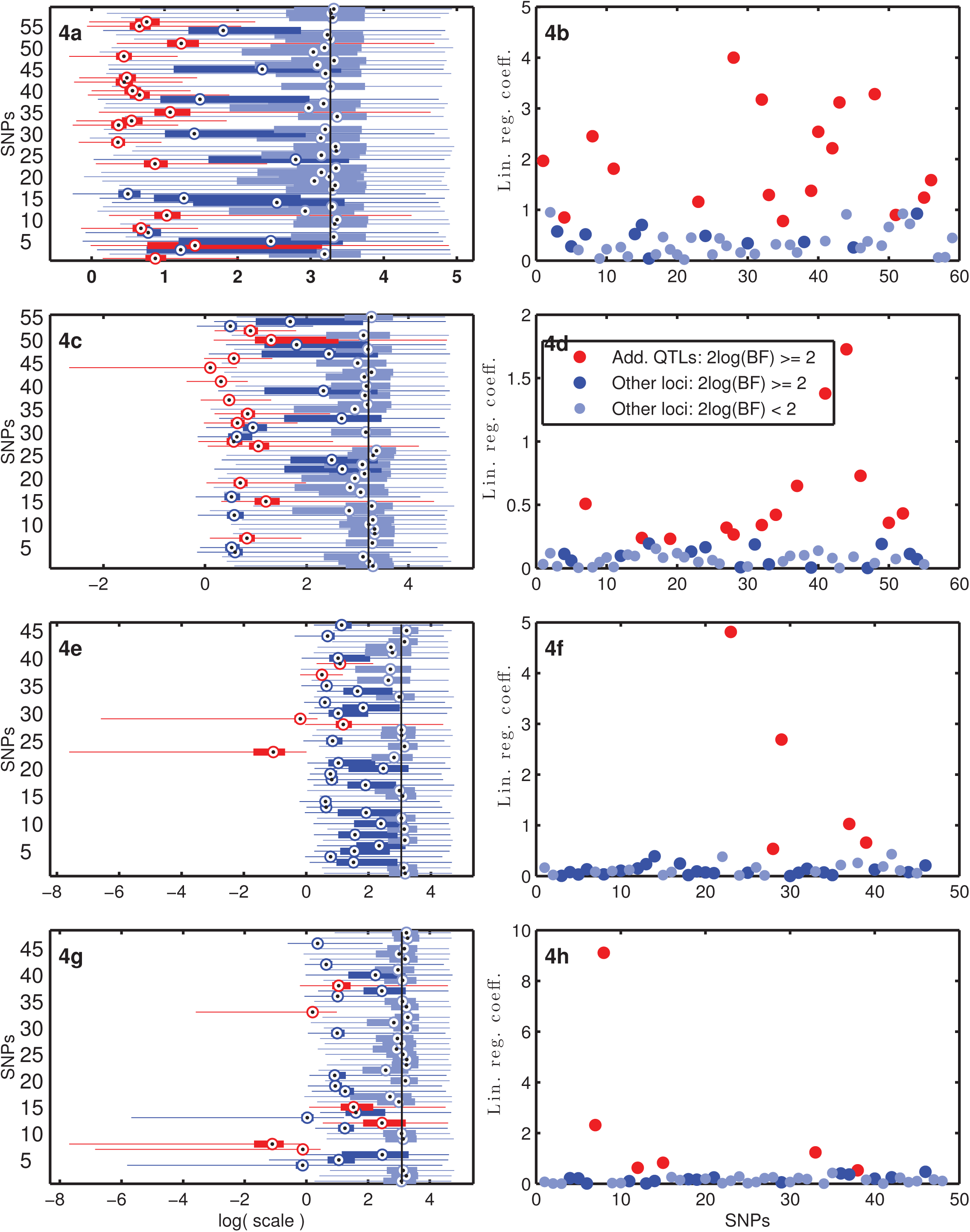
Comparison of loci associated with growth rate variation found by GPR and linear modelling. Left-hand column: box plots summarising the marginal posterior distributions of the scale hyperparameters corresponding to the different SNPs; dots represent the median, edges of the boxes mark the 25^th^ and 75^th^ percentiles, and whiskers indicate the full range of samples from the posterior. The vertical black line denotes the median under the prior. Right hand column: the vertical scale represents the magnitudes of coefficients from an ordinary least squares. multiple linear regression of the same trait against the same SNPs. Each row corresponds to a different growth condition. From top to bottom, these are: Zeocin (Figs. 3a and 3b), Lactose (Figs. 3c and 3d), Maltose (Figs. 3e and 3f), and Cadmium Chloride (Figs. 3g and 3h). Colours denote evidence of association. Red: additive QTLs for which GPR also found positive evidence of association (Methods); dark blue: loci for which only GPR found positive evidence of association; light blue: loci for which no evidence of association was indicated by either approach.

Similarly, for Lactose, GPR explained all of the missing heritability using a combination of additive QTLs and additional loci. For four of these, Chr16: 669064, Chr2: 36827, Chr4: 1416346 and Chr1: 202945 (loci 53, 5, 16 and 4 respectively in **Figs. 3c** and **3d**) evidence of association (Pr(scale ≤ 10) > 0.94 and (2log(BF) ≥ 10] was as strong as that for most of the additive QTLs. None were in strong LD with additive QTLs; the strongest pairwise correlation between one of these loci and an additive QTL was −0.095.

Maltose was the only growth condition for which a significant pairwise interaction accounted for a substantial proportion of missing heritability^8^. GPR accounted for all of the missing heritability for Maltose while confirming the importance of these two loci on chromosomes 7 and 11 (**Figs. 3e** and **3f**).

Our results for the other yeast growth conditions (**Supplementary File 2, Figs. 6 – 17**) suggest that they often followed a similar pattern to that observed for Zeocin and Lactose. This could even be the case when missing heritability was modest. This is illustrated by results for growth in conditions incorporating Cadmium Chloride, for which *H*^2^ ≈ 0.96. Although an additive model incorporating 6 loci apparently accounted for of phenotypic variance (**Supplementary File 1**), GPR found positive evidence of association (2log(BF) ≥2) for an additional 14 loci. For two of these, evidence of association (Pr(scale ≤ 10) > 0.99 was very strong. However, once again, at least one, Chr4: 696694 (locus 13 in **Figs. 3g** and **3h**), was effectively invisible to additive modeling (nominal univariate p-value = 0.2186 and effect size ranked 43^rd^ out of 48 in a multiple linear regression model).

### Statistical Power - the value of biological replicates

Although it is common to use averages of phenotype measurements from biological replicates, we would expect GPR to gain power by using all such measurements. This is because of the strong information they provide concerning plausible values for the unexplained variance. To investigate this, for selected yeast growth conditions we constructed pairs of new datasets by subsampling the existing data. Only one of each pair had replicates, but both consisted of the same number of phenotype measurements. Consequently differences in inferences based on these datasets must lie in the presence of replicates rather than the amount of data. Comparison of heritability estimates (**Figure 4**) indicated that the power to estimate non-additive variance was significantly increased when replicates were used. Estimates of additive variance were less variable.

**Figure 4.**
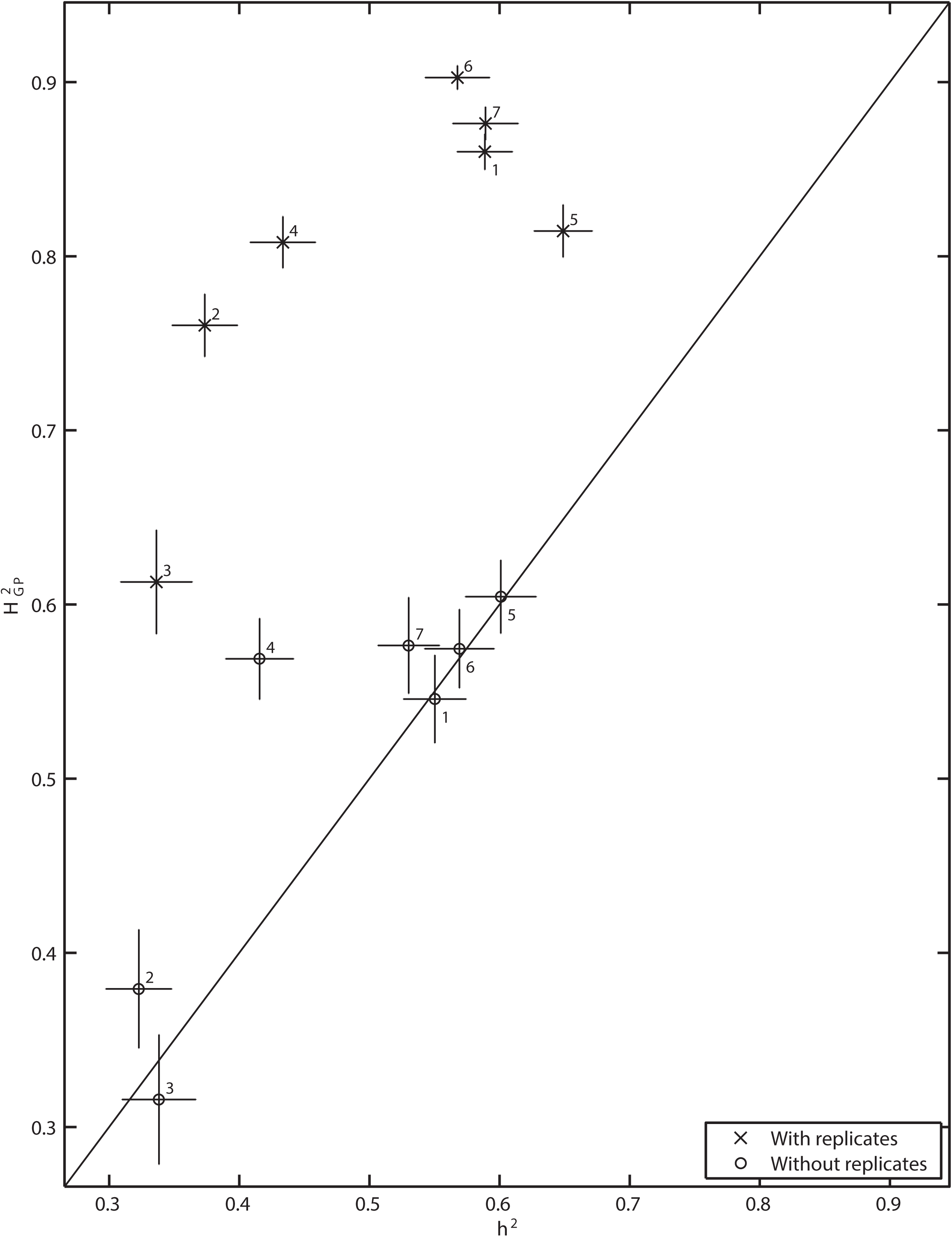
Effect of biological replicates on estimation of non-additive variance. Estimates of explained variance were made using pairs of datasets for 7 different conditions. One of each pair included replicate measurements for some strains. The other incorporated only a single measurement per strain, but included more strains so that both datasets consisted of the same number of samples. Vertical axis: *H*^2^_GP_. Horizontal axis: Multiple linear regression estimates of - *h*^2^ using the same subset of SNPs. Each point corresponds to a different growth condition. Circles indicate estimates based on only a single phenotype measurement for each strain. Crosses denote estimates based on using replicate measurements. Numbers indicate the condition corresponding to a point: 1 – Cobalt Chloride, 2 – Formamide, 3 – Galactose, 4 – Indoleacetic Acid, 5 – Lactate, 6 - YPD:4C, 7 – Zeocin. Comparison of heritability estimates indicates that the power to explain non-additive variance is greatly increased when replicates are used. In contrast, estimates of additive variance are less variable. Vertical error bars denote around the mean GPR estimates. Horizontal error bars indicate the standard error in the linear regression estimate (Methods). For finite sample size, the frequentist and Bayesian estimates of uncertainty, as indicated by the horizontal and vertical error bars, cannot be directly compared. However, they exhibit similar scaling properties with sample size: asymptotically, Bayesian marginal posterior distributions tend to normality with a standard deviation that scales inversely with the square root of the sample size. The solid line is simply a guide to the eye indicating where estimates are equal.

To provide further insight into the effect of using phenotype measurements from replicates, we constructed a simple, toy model. Each output, y_i_, was generated from a simple non-linear function of a single input variable, x_i_, with added Gaussian noise (**Methods and Supplementary File 2 Table 2 &** **Figure 5**). Small numbers of samples generated from this model could not capture all significant variation in the function. Application of GPR demonstrated that the resulting uncertainty in inference of the underlying function induced a multimodal posterior distribution: plausible explanations of the data accounted for observed variation in y either as predominantly owing to variations in x or as noise. However, measurements of replicate data altered the proportion of probability mass in these modes by providing more information about the probable level of noise (**Supplementary File 2, Table 2** and **Figure 5**). Consequently, they not only improved estimation of explained variance but also increased the evidence for the relevance of the input, x.

**Figure 5.**
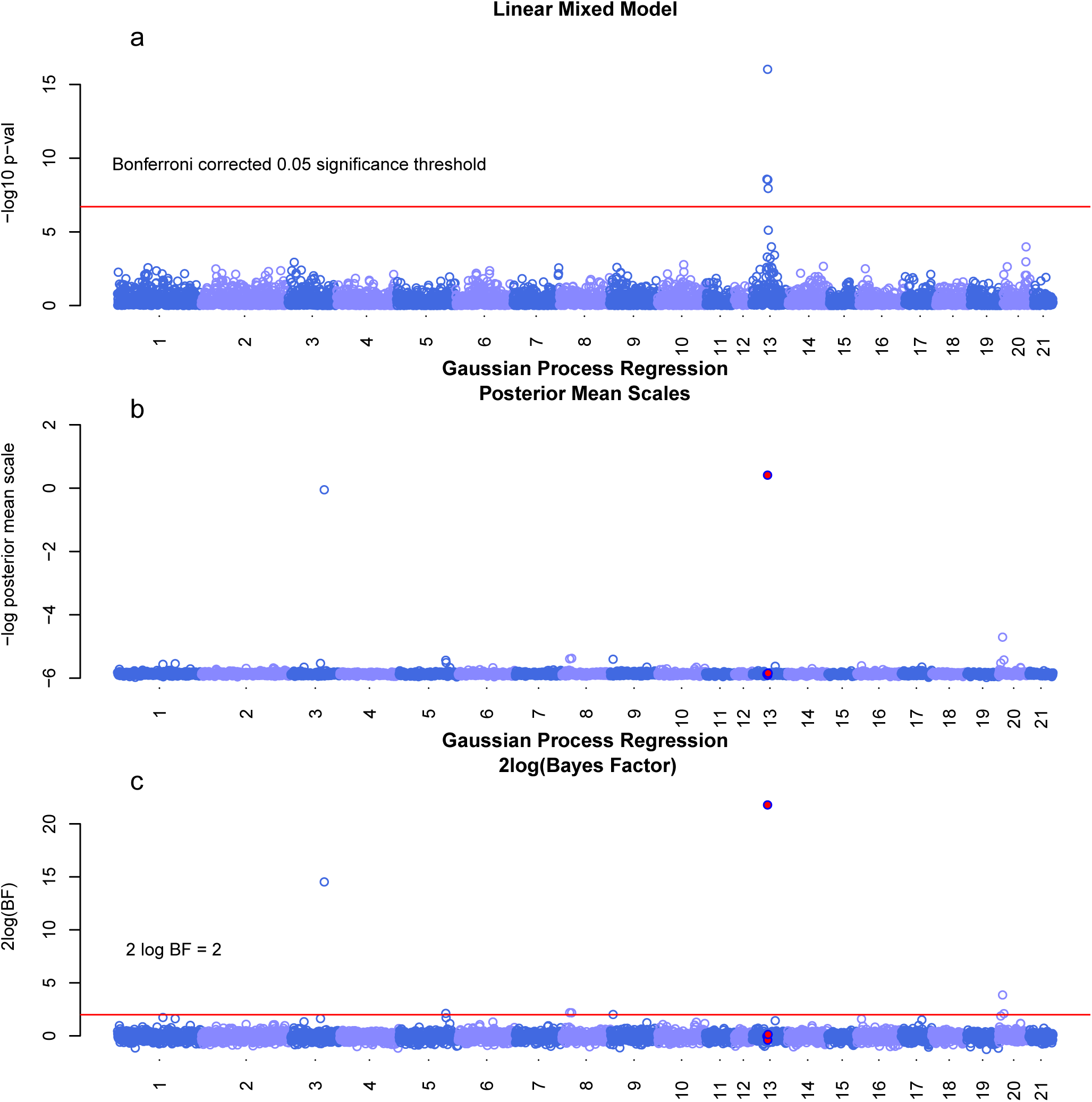
Manhattan plots comparing loci associated with expression of CD45RC in CD8 cells found by GPR and linear mixed model analysis. Horizontal axis: genomic position of pruned set of 5736 SNPs; numbers correspond to chromsomes. **a** Vertical axis: -log_10_ p values from standard univariate association tests using a Linear Mixed Model (LMM) as implemented in GEMMA^27^; horizontal red line indicates the Bonferroni corrected threshold of significance at the 0.05 level based on the unpruned set of 262,052 SNPs. **b** Vertical axis:-log10 posterior mean of the scale hyperparameters used to indicate relevance in the GPR model; smaller posterior mean scales (larger values of-log10 posterior mean scales) imply stronger association. Red filled circles indicate marginally significant SNPs (after Bonferroni correction). **c** Vertical axis: 2log(Bayes Factor). No absolute level of significance can be ascribed to the posterior means of the scales, but we also compute approximate marginal Bayes Factors (**methods**). The red horizontal line corresponds to a value of 2. Values of 2, 6 and 10 for twice the natural logarithm of the Bayes Factor are commonly taken to indicate positive, strong and very strong evidence respectively^25^.

### Expression of CD45RC in CD8 cells in Heterogeneous Stock Rats (Rattus Norvegicus)

To indicate the wider potential applicability of our GPR method, we applied it to an immunological phenotype (expression of CD45RC in CD8 cells) measured in heterogeneous stock (HS) rats^26^.

Unlike the yeast data, the possibility of confounding effects due to relatedness and unmeasured environmental covariates could not be ruled out in this case. To account for this, we applied an enhanced version of our GPR model including an additional term equivalent to a random effect in a linear mixed model (**Methods**).

As this population of HS rats also exhibits substantial long-range LD, we again pruned to remove highly correlated SNPs but retained independent SNPs with significant marginal associations (**Methods**). This reduced 262,052 high-quality genotyped SNPs to 5,736 approximately independent SNPs for 540 individuals.

A standard Linear Mixed Model (LMM) analysis^27^ (**Figure 5a**) shows a single strong marginal hit on chromosome 13 with lead SNP, Rn34_13050905766 (p value 4.5e^-17^,MAF=0.19). However, GPR (**Figure 5b & 5c** **& Supplementary Table 3**) finds strong evidence for the importance of Rn34_13050905766 and an additional SNP on chromosome 3, Rn34_3134037254 (marginal p value 2.2e^-2^, MAF=0.44). There is also weaker evidence at other loci, indicated by bimodal posterior distributions for the hyperparameters indicating relevance (**Supplementary figure 18**). The strongest of these secondary signals was for Rn34_20009896057 (marginal p value 4.1e^-3^, MAF=0.40).

GPR gains power to identify associated SNPs by averaging over probable models, but does not identify the most probable models themselves. Nevertheless, given a small set of SNPs identified by GPR, we may probe the implied models. **Supplementary Figure 19** indicates a dominance model for Chr3: Rn34_3134037254 (where at least one copy of the minor allele results in lower CD45RC expression), while **Supplementary Figure 20** suggests interactions between chr3: Rn34_3134037254 and both chr13: Rn34_13050905766 and chr20: Rn34_20009896057 where expression increases with allele dosage of chr13: Rn34_13050905766 and chr20: Rn34_20009896057, but at levels controlled by the presence of at least one minor allele of chr3: Rn34_3134037254.

Classical likelihood ratio tests to compare the fit of different models (**Methods**) support these conclusions (**Supplementary Table 4**). While the maximum likelihood estimate (MLE) of the additive effect of Rn34_13050905766 alone explains 13.7% of heritability, a model incorporating, additionally, the additive effect of Rn34_3134037254 together with the interaction between the two provides a better fit (p value 6.2e^-05^), explaining an additional 4.5%. A more complex model involving all three SNPs and incorporating both dominance and interaction terms also provided a clearly significantly better fit than a null model consisting of only the additive effect of chr13: Rn34_13050905766 (p value 9.0e-^11^) and explained 26.8% of heritability. This MLE is consistent with a conservative GPR estimate of 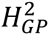 using a model incorporating only these three SNPs (**Online methods, Supplementary Figure 27**). No independent estimates of *H*^2^ were available, but an LMM estimate under the null model gave *h*^2^ = 36.8%. GPR apportions 25% of heritability to a model incorporating the three SNPS and 21% to the random effect, thus increasing estimates by ∼9% for a total of 46%.

Chr13: Rn34_13050905766 (59758577 bp) is in very high LD with a number of SNPs at the beginning of the gene (60,094,038 - 60,205,773 bp) coding for the Ptprc (CD45) enzyme. It has a Pearson correlation of 0.997336 with 3 SNPs within this gene for which it was acting as a tag in the GPR analysis (**supplementary figure 25**). CD45 is a tyrosine phosphatase which undergoes alternative splicing of three variable exons to yield multiple isoforms of which CD45RC is one.

Chr3: Rn34_3134037254 lies within the Slc24a3 gene (145,760,074 −146,259,491) and is in high LD with many other SNPs in this gene (**supplementary figure 24**). Slc24a3 encodes the solute carrier family 24 member 3 protein (also known as NCKX3) that plays a critical role in the exchange of calcium, potassium and sodium ions into and out of the cell.

A link between the expression or activity of Slc24a3 and expression of CD45RC might be provided by the regulatory effect of two other molecules: CD86 and CD28. CD86 and CD28 are co-stimulatory molecules that interact to affect activation of naive T cells and their subsequent differentiation^28^; inhibition of CD86 signaling is one pharmaceutical strategy used to counteract inflammatory disease such as rheumatoid arthritis^29^. However, increased expression and activity of Ca++/Na+ exchangers has also been shown to lead to decreased expression of CD86^30^. This might be expected to have a general immunosuppressive effect^30^. However, it might also have an effect specific to expression of particular CD45 isoforms as it has been shown that CD28 affects expression of hnRNPLL^31^, a trans acting factor that regulates exon silencing of the variable exons of Ptprc mRNA in CD4 and CD8 cells^32^.

## DISCUSSION

To demonstrate proof-of-principle, we have analysed an existing dataset of 46 yeast traits and a rat gene expression phenotype. For the yeast phenotypes, we have shown that GPR can explain much of the known missing heritability using SNPs. It does so even in the presence of high order interactions possibly involving ∼20 loci when some SNPs display only weak marginal effects and significant pairwise effects are absent. In the rat phenotype, we uncovered strong evidence of at least one novel interaction together with indications of a model incorporating three SNPs (only one of which had a significant marginal association) that explained significant additional heritability.

The significant loci GPR identifies typically include most independent, additive QTLs, but also a number of additional loci not in strong LD with any additive QTL. In some cases, such as Chr4: 832287, Chr16: 669064, and Chr4: 696694 for yeast growth in the presence of Zeocin, Lactose and Cadmium Chloride respectively, these additional loci were found to be at least as important as the additive QTLs in determining the phenotype but exhibited no marginal effects. Combined with the reported absence of significant pairwise effects for these growth conditions^8^ and the demonstration that GPR is not overfitting (**Fig. 2**), this provided clear evidence that GPR can identify loci that exert strong effects on the phenotype only through higher order interactions. Such loci would have been challenging to identify using methods that employ a filter based on low order effect sizes.

Our findings are consistent with the suggestion that much additive variance is, in fact, an artifact of non-additive interactions^6,33^. Even, as in the case of Cadmium Chloride, when narrow-sense heritability is very high, GPR identifies a number of important loci with no marginal significance implying a more complex underlying model. High narrow sense heritability does not necessarily imply independent effects.

Lower estimates of *H*^2^ were accompanied by an apparent decrease in the ability of GPR to explain all missing heritability. At the same time, GPR heritability estimates became more uncertain. An interpretation of these results is suggested by clear evidence of two modes in the posterior distribution of 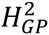 for some conditions (**Supplementary File 2**, **Fig. 3**): as the unexplained variance increases models of the data which explain more variance as noise become increasingly more probable. Posterior modes corresponding to models which explain all of the missing heritability may continue to be present, but will contain less probability mass. Such modes will become progressively less influential in computing explained variance and the importance of SNPs unless evidence for the models they represent is strengthened by increasing the sample size. Similar issues of weak identifiability of the variance components when heritability is lower have previously been noted in Bayesian models for estimation of additive and dominance components of genetic variance^34^.

The strength of the sparsity prior can also affect heritability estimates. The prior should favour models comprising fewer SNPs. We used two different approaches to specifying this prior, each with its own advantages (**Methods**). The first approach, while conservative, might be too stringent for highly polygenic traits resulting in underestimates of *H*^2^. It is likely that this is why GPR heritability estimates for yeast growth rates were lower in four of the growth conditions than linear mixed model estimates of narrow-sense heritability based on all SNPs (**Results**). For these traits, a highly polygenic architecture seems likely: broad-sense and narrow-sense heritability were almost equal while additive QTLs alone explained only a portion of heritability. In general, if such information about the likely genetic architecture were available (for example from an analysis of independent data) this could be used to guide the choice of prior. Otherwise, we would recommend use of a more flexible, hierarchical prior as we used for the analysis of the rat data (**Methods**), which can increase power in the case of more polygenic traits.

There have been a number of reports of interactions in human disease and quantitative traits^20,21,35-37^. However, the overall contribution of these to heritability is still difficult to determine. The two key challenges are computational tractability and statistical power. Our results have implications for both.

The computational cost of exhaustive search grows exponentially with the order of interaction considered. Consequently, such approaches are unlikely to be tractable for higher-order interactions. For example, a recent exhaustive search for pairwise interactions in the WTCCC data^20^ required a total computing time of 950 compute years. Thresholding on lower-order effects can reduce the scale of the search but may also sacrifice considerable power; we have identified a number of important loci without marginal effects that are apparently also not even involved in detectable pairwise interactions. In contrast, the computational cost of the GPR sampling algorithm does not depend directly on the order of interactions. Nevertheless, it is computationally intensive. Our current implementation, using a Graphics Processing Unit (GPU), enables application to significantly larger datasets than we have reported in this paper (**Methods**), but we anticipate that significant further improvements can be made. One idea we are pursuing is to exploit the sparse nature of our model. Currently, most of the computations performed to generate the proposed next state for our MCMC sampler make negligible contributions: at any given iteration, the vast majority of SNPs are, effectively, not included in the current model. These calculations can be replaced by a single approximate computation, while retaining the validity of the overall algorithm.

Even when computationally feasible, approaches that explicitly test specified interactions require very large sample sizes to maintain power in the face of the additional multiple testing burden. GPR gains power by averaging over possible interaction models rather than identifying the specific partners in the interaction. In addition, we have observed that power is further substantially improved by using phenotype measurements from replicates. This additional power derives from the strong information such replicates provide concerning plausible values for the unexplained variance. In contrast, with much less freedom to adapt to additional data, linear models derive little added value from replicates.

It is important to note that the nature of the replicates does not affect this conclusion. It applies whenever one might reasonably posit a model similar to the one used here (**Methods, equation (1)**); power might be gained from replicates in a broad range of settings ranging from phenotypes of genetically identical individuals to replicate measurements of cellular traits based on averages over many genetically identical cells such as gene expression levels. Consequently, even though our study was limited to yeast, our findings have important practical implications for study design. Firstly, cohorts consisting of a significant number of monozygotic twins, such as the TwinsUK cohort (http://www.twinsuk.ac.uk), might offer a way to estimate the contribution of non-additive effects to quantitative traits and disease in humans while increasing power to identify important interacting SNPs with weak marginal effects. Secondly, generating replicates in cellular phenotypes, such as transcriptome studies, combined with analysis by GPR may be useful to delineate the role of interactions in gene regulation.

There are a number of natural extensions to this work. A straightforward extension of the GPR model can allow for the possibility of interactions between SNPs and measured covariates. We are currently exploring this idea to search for evidence of gene-environment interactions in data from the UKBiobank (http://www.ukbiobank.ac.uk/). An interesting additional avenue of research would involve an extension to analyze multiple traits. Building on other recent work^38^ a multi-output Gaussian Process framework could be employed to search for evidence of gene-gene and/or gene-environment interactions with the potentially greater power provided by correlated phenotypes.

## METHODS

### Code Availability

An implementation of our GPMM method which can run on CUDA enabled Graphics Processing Units will be made freely available for academic use at http://www.stats.ox.ac.uk/∼sharp/

### Yeast Data Preprocessing

The yeast genotype data^8^ comprised 11,623 SNPs for a panel of 1008 prototrophic, haploid segregants, constructed from a cross between a laboratory strain and a wine strain. Phenotypes consisted of replicate measurements of growth rates on 46 different media. To minimize the number of essentially redundant explanations of the data, we pruned the single nucleotide polymorphisms (SNPs) so that no pair had a correlation of > 0.8. However, SNPs already identified as quantitative trait loci (QTLs) under an additive model were always retained. This procedure reduced the number of candidate loci drastically to ∼ 60 so that each chromosome was represented, on average, by just 3-4 SNPs.

### Rat Data Preprocessing

The rat genotype data comprised 262,052 high-quality genotyped SNPs [ref amelie] for 540 heterogeneous stock (HS) rats. The phenotype consisted of expression levels of CD45RC measured in CD8 cells. We obtained the data from https://www.ebi.ac.uk/arrayexpress/experiments/E-MTAB-2332/files/.

As this population of HS rats also exhibits substantial long-range LD, we again pruned to remove highly correlated SNPs so that no pair had a correlation of > 0.8. In this case, there was a single eQTL on chromosome 13 for which a number of SNPs in high LD showed significant marginal signals of association after Bonferroni correction. To ensure that we retained those SNPs with possibly independent signals we used a greedy approach: firstly we pruned all SNPs with a pairwise correlation of > 0.8 with the lead SNP. From the SNPs that survived the first step we identified the SNP with the second strongest marginal association and pruned again based on correlations with this SNP. We repeated this process for successively less strongly associated SNPs until all such SNPs had been considered. Finally, we performed one further round of pruning to obtain a set of 5,736 approximately independent SNPs.

In contrast to the yeast there was a significant possibility of confounding effects together with measurements for suspected relevant covariates: sex and batch. To account for the latter we simply created a new phenotype from the residuals of the multiple regression of the original phenotype on these covariates. To allow for unmeasured confounding effects, we incorporated an additional term, analogous to a random effect, into our GPR model as described below (**Gaussian Process Regression**)

### Gaussian Process Regression

We assume that, for the *i*^*th*^ of N, haploid individuals, we have observed a quantitative trait value, *y*_*i*_, and a genotype vector, 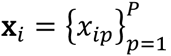, of allele dosages at *P* markers where each *x*_*ip*_ ∈ {0,1} is the dosage at the *p^th^* marker. The assumed generative model has the following form:

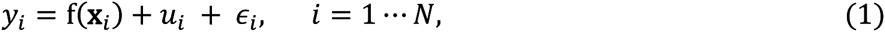

where f(**x**_*i*_) is an unknown, arbitrary function of the allele dosages, *u*_*i*_ is equivalent to the random effect in a standard mixed effects model and 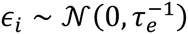 represents additive Gaussian noise of precision *τ*_*e*_.

The second term is commonly used in association analyses to account for potential confounding factors due to unmeasured environmental effects that correlate with genotype. The vector of random effects for all individuals has covariance structure determined by the relatedness of the individuals 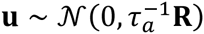. For our analysis of the yeast data we did not incorporate such a term as the experimental design precluded the possibility of such effects: all strains shared the same environment. However, for the analysis of the rat phenotype we did incorporate this term. **R** was the centred, scaled, realised relationship matrix computed from SNPs estimated using functionality in GEMMA^27^.

No explicit, parametric form is assumed for **f(x_*i*_)**, which permits great flexibility in modeling interactions of unknown order and effect size. Instead, a Gaussian Process prior (𝒢𝒫) is placed over the functions themselves. This prior imposes constraints by making assumptions about properties of plausible functions such as their smoothness. To achieve this, the defines a probabilistic relationship between the genotypes for the *N* individuals and the corresponding *N-*dimensional vector of regression function values, **f**. A priori plausible examples of **f** are modeled as draws from a multivariate Gaussian distribution:

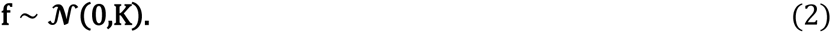

While we may draw many different vectors of values from this distribution, their values will be constrained by the *N* × *N* covariance matrix **K**. This is the crucial element of this prior.

### Covariance function

The elements of **K** are generated as functions of the regressors, **x_*i*_**, in a way that reflects prior beliefs about properties of plausible functions. A very general assumption underpinning any data-driven estimation method is that similar inputs result in similar outputs. In this context, the notion of similarity between genotypes is defined by a covariance function, k(**x_*i*_,x_*j*_**), which generates the elements, K_*ij*_ of **K**.

The covariance function embodies important prior assumptions about the regression function. A wide choice is available. For example, it is perfectly possible within this framework to use a kernel that permits only linear functions. However, we wish to allow for the possibility of highly non-linear interaction effects. A covariance function that permits a very broad class of such functions is the squared exponential covariance:

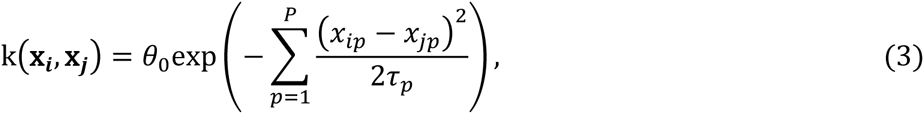
where and *θ*_*0*_ and *τ*_*p*_ ∈ {*τ*_*1*_,‥., *τ*_*P*_} are hyperparameters of the model. This choice of covariance function also implies other assumptions about the properties of the unknown regression function. It is possible that other choices might be more appropriate in some situations^22^.

*θ*_*0*_ determines the overall variance in the observed outputs, **y**, due to the regression function, **f**. The *τ*_*p*_ determine characteristic scales in the respective predictor variables. A large value for *τ*_*p*_ relative to the observed range of variation of (*x*_*ip*_ – *x*_*jp*_) indicates that variations in the *p*^*th*^ dimension contribute very little to variations in f; a smaller value indicates the reverse. Consequently, each scale hyperparameter provides a measure of the relevance of the corresponding predictor in explaining variations in the response variable^39^. Therefore, when the *x*_*ip*_ represent allele dosages, we may discover which SNPs are most likely to be associated with the quantitative trait by inferring values for the corresponding scale hyperparameters, *τ*_*p*_.

### Hyperparameter Priors

A priori, we expect that only a small number (if any) of the candidate loci are likely to be associated with variations in the trait. To reflect this, we place a sparsity-inducing prior over the *τ*_*p*_ to ensure a very low prior probability of relevance for any individual SNP. A standard choice is a gamma distribution^40^ which we parameterise in terms of its mean, *μ*_*x*_, and shape parameter, *α*_*x*_:

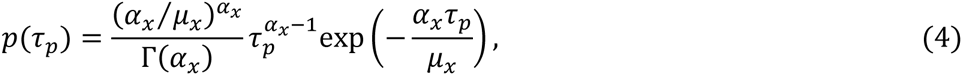
where Γ(.) is the gamma function. A priori, we assume that the *τ*_*p*_ are identically and independently distributed; we choose *α*_*x*_ = 0.5 to ensure that the prior on *α*_*x*_ has no mode and *μ*_*x*_ to incorporate a prior belief that only a subset of SNPs will have significant effects.

A simple way to achieve this is to fix *μ*_*x*_, so that, in *P* draws from the prior, no more than a certain small number, *S*, are expected to have a scale smaller than a given low threshold, *τ*_*r*_. In general, choosing S < 10 and 1 ≤ *τ*_*r*_ ≤ 10 will ensure that any given marker has a very low prior probability of being relevant. In all of the analysis on yeast data we set *S* = 1 and *τ*_*r*_ = 2.

An alternative is to specify a prior distribution on *μ*_*x*_. We again use a gamma distribution:

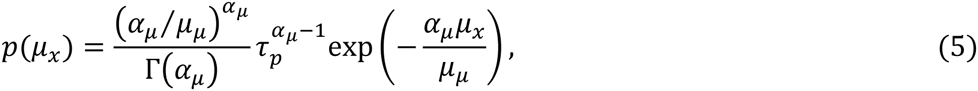

We choose *α*_*μ*_ = 0.5 (again to ensure that this prior has no mode) and *μ_μ_* ≫ 1 to encode a prior belief in a sparser model.

Each approach has advantages and disadvantages. Fixing *μ_x_* avoids one potential source of mixing problems in the Markov Chain Monte Carlo (MCMC) algorithm that we use to perform inference. However, posterior inferences will have some sensitivity to this choice. While the conservative choice described above offers protection from false positive associations, it can also reduce power. For the analysis of the yeast data set we chose to fix *μ_x_* because the number of SNPs was small and strong information was provided by the presence of biological replicates (**Results** and **Supplementary File 2 section 1.6**). However, for the analysis of the rat data, we used the hierarchical prior. We set *μ*_*μ*_ = 5*e*^5^ which implied an initial distribution for each *τ*_*p*_ that was less restrictive than a prior with *μ*_*x*_ fixed as described above. We found this helped with mixing while not precluding inference of a much sparser posterior distribution for the *τ*_*p*_ if this was supported by the data. For example. for the rat phenotype, the posterior distribution of *μ*_*x*_ implied a much sparser distribution for the *τ*_*p*_ than our initialization.

The final elements of the model are the remaining covariance hyperparameter, *θ*_*0*_, the scale parameter, *τ*_*a*_. for the random effect and the noise precision *τ*_*e*_. We place gamma priors over 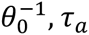 and *τ*_*e*_ with hyperparameters *α*_*θ*_, *μ*_*θ*_, *α*_*a*_, *μ*_*a*_ and *α*_*e*_, *μ*_*e*_ respectively. To ensure a diffuse prior with no mode, we set *α*_*θ*_ = *α*_*a*_ *α_a_* = 0.5. Finally, we set *μ*_*θ*_, *μ*_*a*_ and *μ*_*e*_ to the reciprocal of the observed trait variance. This choice is reasonable given that all plausible models of the data must account for variance on this scale. Note that this prior is uninformative with respect to how the total variance is partitioned between genetic factors (determined by *θ*_*0*_) and non-genetic factors modeled here as Gaussian noise (determined by *τ*_*e*_).

### Relationship to Previous Work

A previous study^41^ proposed a similar use of GPR for identifying QTLs in the presence of epistasis. There are a number of differences between their approach and ours. Two concern the model. Firstly, we incorporate a random effects term. Secondly they employed a more complex sparsity-inducing prior over the *τ*_*p*_: a mixture prior involving additional binary auxiliary variables to indicate relevance. This appears to provide a direct probability of relevance for each locus rather than the more indirect measure provided by the scale hyperparameters. However, this scheme still involves prior specification of which scales imply relevance. Consequently, we do not believe it offers any advantage over our approach. Furthermore, inference for the additional indicator variables necessitates the use of an additional (single-site) Gibbs sampling step. This is known to result in much less efficient sampling when variables are highly correlated^42^. Such a situation might be quite common in this context when there are multiple plausible models of the data not all incorporating the same subset of loci. Consequently, we expect our simpler approach to be significantly more efficient. Additional differences in our inference algorithm and implementation include further steps to improve the mixing efficiency of our Markov chain, speed per iteration, and hence scaleability: we use a number of leapfrog steps to generate each proposal rather than a single step; we adapt the step size and we use a modified mass matrix (**Methods**: **Inference – Hybrid Monte Carlo**). Finally, we have a CUDA implementation of our method that uses a graphics processing unit (GPU) to exploit the significant opportunities for parallelisation of the algorithm.

### Statistical power - Marginalisation of the latent function

Finding evidence of higher order interactions poses significant challenges owing to the curse of dimensionality. On average, linear modeling indicates ∼13 QTL per trait for the yeast phenotypes that we consider^8^. Under the reasonable assumption that interaction models might involve additional loci, we are faced with the apparently hopeless task of attempting to learn a function over, say, 2^20^ combinations of variants with only, perhaps, 1000 samples (N=1000). Nevertheless, it is possible to make inferences about not only the heritability but also the relative importance of individual loci within the interaction model, although it is not possible to identify the nature of the interactions; the method can identify that loci are important for explaining the phenotype, possibly through high order interactions with each other, but does not identify which loci interact with which others.

Two aspects of the model increase power to make inferences in this context. One is the sparsity assumption already described. The second is the fact that we do not need to learn the latent function at all for this purpose. Within this model we can effectively average over plausible functions, by integrating over **f**, but still make useful inferences based on the posterior distribution of the hyperparameters. The hyperparameter posterior contains all the information necessary to identify relevant loci and estimate heritability. The fact that integration over **f** can be done analytically leads to significant computational advantages as, in general, hyperparameters and function values are highly correlated.

After marginalising over **f,** the resulting posterior may be summarised as^22^:

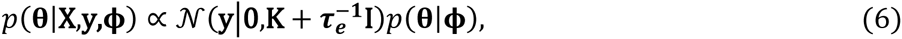
where **X** represents an *N* × *P* matrix of observed genotypes, **θ** represents the set of hyperparameters, 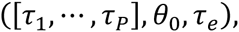 and *p*(**θ|ϕ**) represents the joint prior over these hyperparameters (a product of gamma distributions) parameterised by a further set of fixed hyperparameters, (*α*_*x*_, *μ*_*x*_, *α*_*θ*_, *μ*_*θ*_, *α_e_* and *μ*_*e*_), collectively denoted by **ϕ**.

### Inference – Hybrid Monte Carlo

The posterior defined by (6) is intractable. Nevertheless, analytical expressions for the gradient of the log posterior are readily derived. This permits the application of an efficient form of Markov Chain Monte Carlo sampling algorithm termed Hybrid Monte Carlo (HMC)^23^. The application of HMC to Gaussian Process regression models has been comprehensively described elsewhere^40^. Briefly, the method can be envisaged as simulation of a physical system. The logarithm of the hyperparameters, defines the position, **q**, of a notional particle, (**q** = log **θ**). The logarithm of the posterior over **q**, – log *p*(**q|X, y, ϕ**), defines a potential energy. A vector of auxiliary momentum variables, **p**, are introduced, one element for each dimension. These may be sampled from some distribution independent of **q**:

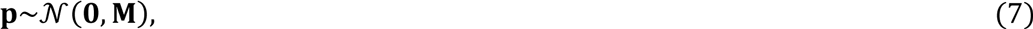

These define a kinetic energy, 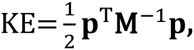 where **M** is a ‘mass’ matrix whose elements can be tuned to improve mixing as described below.

The joint distribution for the system is obtained from the Hamiltonian, 𝓗(**q, p**):

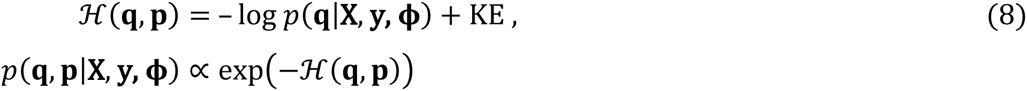

As the joint distribution defined by (7) factorizes between **q** and **p**, any valid sample immediately yields a sample from the posterior over (**θ**) To sample from (7), a new state, (**q′, p′**) is proposed from the current state, (**q, p**), by simulating a trajectory for the particle under Hamiltonian dynamics, a system of differential equations. Under such a trajectory, 𝓗(**q,p**) is, in principle, conserved. However, in practice, discretization of the system of differential equations introduces errors. Nevertheless, discretization schemes can be devised which are both reversible and volume preserving. Consequently, it can be shown that acceptance of the proposal based on a simple Metropolis-Hastings acceptance test leaves *p*(**q, p|X, y, ϕ**) invariant.

The effect of the momentum variables is to improve on the slow random walk exploration of ordinary MCMC by enabling more distant proposals with a good chance of acceptance. Nevertheless, the acceptance probability can be sensitive both to the step size, *ε*, used for discretization and, to a lesser extent, the length of the trajectory, *L*. In particular, the optimal *ε* can vary in different regions of the state space. To address this, we resampled *ε* for each trajectory:

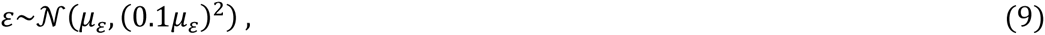
where *μ*_*ε*_ is initialized as 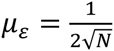 and then adapted during a burn in period so as to maintain an acceptance rate of between 0.6 and 0.7.

Avoidance of random walks and reduction in the autocorrelation of the chain also depends on *L*. However, as the computation of each step is expensive, it is wasteful to compute a proposal based on a long trajectory that is subsequently rejected. We chose a compromise: we used a short trajectory length (*L* = 10) and used partial momentum refreshment^43^; instead of resampling **p** according to (6), we updated it partially:

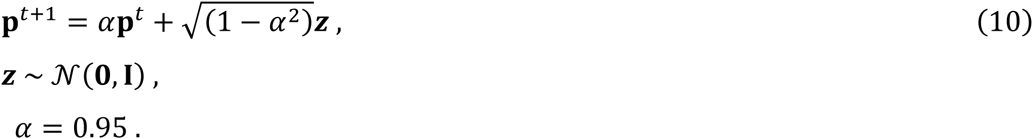

An additional issue is that optimal step sizes might vary across dimensions owing to differences in the natural scales of the different variables and the underlying geometry of posterior. In general, this can be addressed through tuning of the mass matrix, **M**, in (7). Sophisticated approaches for estimating a full covariance for **M** scale poorly with dimension^44^. We take a simpler approach which addresses only the diagonal of **M**: firstly we use a logarithmic transformation of the hyperparameters which mitigates the different scales over which the different types of hyperparameter naturally vary; secondly, when we use the hierarchical prior over *μ*_*x*_(5), we use constant factors on the diagonal of **M,** proportional to log *μ*_*μ*_ to reduce the step size in the direction of the variance hyperparameters, *μ*_*θ*_, *μ*_*a*_ and *μ*_*e*_ relative to that in the direction of *μ*_*μ*_ and the *τ*_*p*_.

### Diagnosing Convergence

To check that our Markov chains had reached stationarity, for each experiment we ran two different chains randomly initialized from the prior and diagnosed convergence using the potential scale reduction factor (PRSF)^45^ as implemented in the CODA R package. For the yeast phenotypes, when all 95% upper confidence limits on PSRFs for individual hyperparameters were approximately 1.1 or lower, we were satisfied that further decreases in the variance of the approximate posterior of the hyperparameters would not significantly alter conclusions.

Convergence for most hyperparameters was often very fast, requiring computation of fewer than 5000 simulated trajectories (50,000 time steps of the simulated Hamiltonian dynamics). However, for some growth conditions there was considerable evidence of multimodality and convergence required computation of ∼ 35,000 trajectories (350,000 simulation steps). For only one condition, Manganese Sulphate, were regions of significant probability mass apparently so isolated that sampler appeared far from convergence even after 35000 trajectories.

### Scaleability

The computational cost of the GPR sampling algorithm does not depend directly on the order of interactions. Nevertheless, it is computationally intensive. For *N* individuals and *P* markers the computation time of each iteration scales as 𝒪(*N*^*3*^ + *PN*^*2*^). For *P* ≫ *N*, the *PN*^*2*^ cost of recalculating the covariance matrix and the gradients required for computing the MCMC proposal dominates. However, the most expensive steps offer a number of options for parallelization. In particular, those operations leading to the apparent linear dependence on the number of markers may be performed independently. We have developed a CUDA implementation that uses a Graphics Processing Unit (GPU) to exploit these features of the algorithm.

**Supplementary Table 5** shows the variation in CPU time, of a single leapfrog step for values of P in the range 1000-500000 and N=500 and 1000. As explained above (**Inference – Hybrid Monte Carlo**), each MCMC iteration might require a trajectory involving multiple leapfrog steps (we used 10 or 20) although a single step would also be valid. In addition, like any MCMC method, it is difficult to predict how many iterations might be required for convergence in any given case. As an example, for the rat phenotype (N=540,P=5736), we ran each MCMC chain for 30,000 iterations consisting of 20 leapfrog steps each. This took ∼35.4 hours. Nevertheless, we have often found that far fewer iterations are required to achieve approximate convergence. SNPs often fall quite quickly into three categories: strongly associated, weakly associated and unassociated. However, the evidence for weakly associated SNPs tends to be reflected by secondary modes of the posterior, which are visited infrequently by the MCMC sampler, Consequently, much longer runs are required to achieve full convergence for these SNPS (compare **Supplementary Figure 30** to **Supplementary Figures 27 & 28**).

### Estimating heritability

GPR partitions trait variance between the model, noise and, when present, the random effect term. For a very highly polygenic trait, some of the variance attributed to the random effect term might reflect genetic effects not captured by the model, f(**x**). However, we make the assumption that all variance explained by this term is attributable to confounding and that f(**x**) and **u** are uncorrelated. Consequently, in both cases, we can estimate 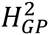 as the fraction of variance in the trait, Var(**y**), explained only by f(**x**).

For single, point estimates 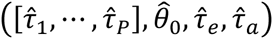 of the hyperparameters, we may estimate the variance components as^46^:

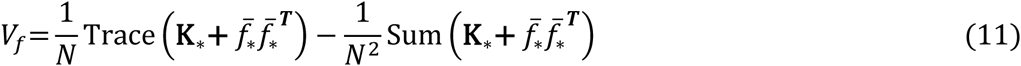

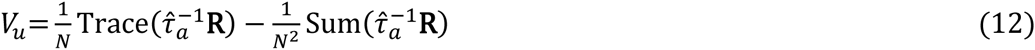

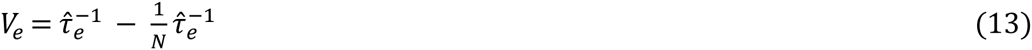

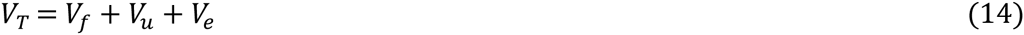
where Sum(**R**) returns the sum of all elements of the matrix 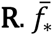 is the posterior mean function (20), and **K**_*\**_ is the covariance of ***f***_*\**_, each element of which is given by (21) using the estimated 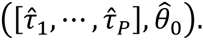 The expected heritability can then be estimated from these variance components as^46^:

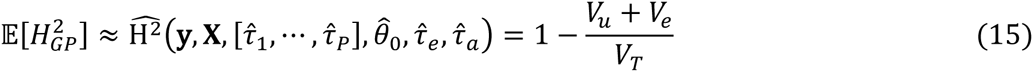
where **X** is the matrix of genotypes and we assume that all variance due to purely genetic factors is captured by *V*_*f*_.

Given a posterior distribution over the hyperparameters we can, in principle, average over the uncertainty in their values to obtain an estimate that is a function of only the observed data 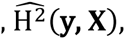 and is robust to overfitting. We approximate this estimate with a Monte Carlo average using the output of our MCMC sampler:

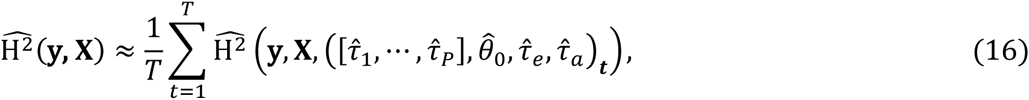
where the sum is over *T* samples from the posterior.

In practice, for the yeast data (for which we did not employ a random effect term), we found that a much simpler computation to estimate 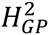 as the fraction of variance in the trait, Var(**y**), not accounted for by noise gave very similar results:

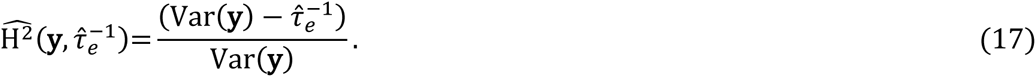

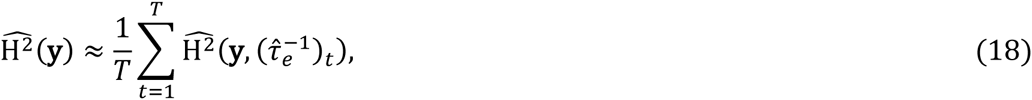

### Identifiability of *V*_*f*_ and *V*_*u*_

When the samples do not contain a significant proportion of replicates, and confounding effects are anticipated, heritability estimation is complicated by non-identifiability of *V*_*f*_ and *V*_*u*_: while using *V*_*f*_ to explain some variance with SNPS of strong effect, GPR can, in principle, also simultaneously use *V*_*f*_ to explain some variance using many other SNPs of small effect. This problem can be mitigated by using a more stringent prior on the relevance of individual SNPs (larger *μ*_*μ*_), but at the possible expense of reduced power to identify truly associated SNPs. An alternative approach is to use two sets of runs. The first set use GPR to identify probable associated SNPs; the second set use *the same prior* but only the subset of probably associated SNPs identified in the first run. We took this latter approach in the analysis of the rat CD45RC expression phenotype. The second run incorporated only the three SNPs found to be most probably associated (**Figure 5b & c**). Posterior distributions of heritability and variance components *V*_*u*_ and *V*_*e*_ thus obtained (**Supplementary figure 27**) were consistent with maximum likelihood estimates of mixed models incorporating additive, dominance and interaction effects for these SNPs (**Supplementary figure 27 and Supplementary Table 4**).

### Independent estimation of broad-sense heritability for yeast phenotypes

Broad-sense heritability was estimated as described in the online methods section of the study that generated the data^8^ using replicated segregant data and a random effects analysis of variance. This involves partitioning variance into a random effect for segregant and a random effect for non-genetic noise. This was implemented using the ‘lmer’ function in the lme4 R package^47^.

### Estimating narrow-sense heritability for yeast phenotypes

Reported estimates of narrow sense heritability from the full set of 11,623 SNPs were computed using a linear mixed model incorporating an estimated relatedness matrix for all pairs of segregants as implemented in the rrBLUP R package^48^. Standard errors were computed using leave-one-out jackknife. Using the same code, we found the leave-one-out jackknife procedure to be very computationally intensive. Instead, we computed estimates for the pruned subset of SNPs by multiple linear regression using the simpler and computationally cheaper method of least squares. We also used leave-one-out jackknife to estimate standard errors and to correct for bias. Comparison of estimates using rrBLUP on the pruned subset of SNPs (but without computing standard errors) indicated that no bias was introduced by this difference in approach (correlation of h^2^ estimates = 0.98).

In both cases, we followed the study that generated the yeast data^8^ and used one randomly chosen measurement of the replicate phenotypes for each segregant. For the subset of SNPs, we found that using all replicate measurements made no difference within the limits of standard error.

### Out of sample prediction

The output of GPR can be used to predict the phenotype, *y*_*\**_, of a previously unseen individual given their genotype, **x**_*\**_. For fixed hyperparameters, the posterior distribution over the unknown, latent function, **f**, induces a Gaussian predictive distribution^22^:

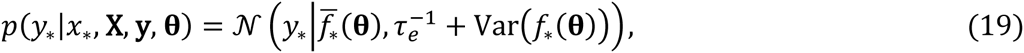

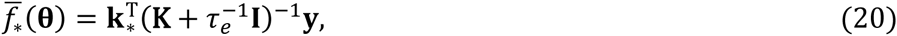

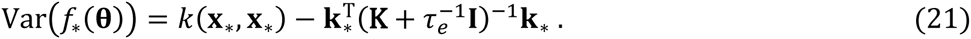
where 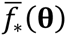 and (*f*_*\**_(**θ**)) are, respectively, the predicted mean and variance of the latent function, *f*_*\**_, corresponding to genotype **x**_*\**_, **k**_*\**_ is the vector of covariances between **x**_*\**_ and the training instances **X**, and *k*(**x**_*\**_, **x**_*\**_) is the prior variance of *f*_*\**_.

To make predictions we average over the values of the hyperparameters with respect to their posterior distribution to obtain a marginal predictive distribution which we approximate with a Monte Carlo average using *T* samples from the posterior:

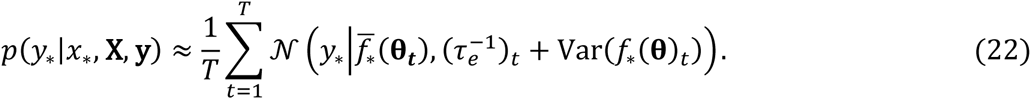

The right hand side of (22) is a mixture of Gaussians. If we assess the quality of predictions using mean squared error, the asymptotically optimal prediction, 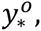 for a given **x**_*\**_ is given by the mean of this distribution. This is obtained simply as the Monte Carlo average of the mean predictions of each Gaussian:

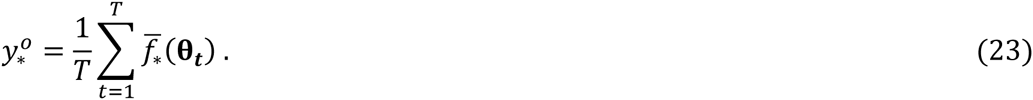

The expected error in this prediction depends on the variance of *y*_*\**_ under *p*(*y*_*\**_|*x*_*\**_, **X, y**). This can be shown to be:

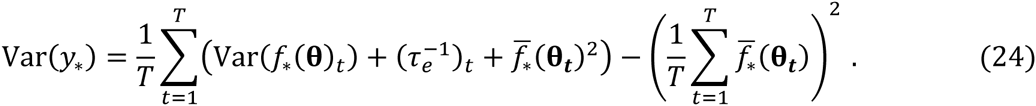

The uncertainty in *y*_*\**_ expressed by the right hand side of (24) is comprised of three different components: uncertainty in the values of the hyperparameters, uncertainty owing to unexplained variance (noise) and uncertainty in *f*_*\**_ for any fixed setting of the hyperparameters. Assuming homoscedastic noise, prediction errors will be greatest when uncertainty in the value of *f*_*\**_ is greatest. This will be the case when **x**_*\**_ is very different from the genotypes of the individuals used for training.

The predictive distribution of (22) comprises the sum of a large number of approximately independent terms. Consequently, the Central Limit Theorem indicates that the mean squared errors in predictions made using (23) should be Gaussian distributed with zero mean and variance given by (24). We verified that this was the case (**Supplementary File 2**, **Figure 4**)

### SMSE

As a measure of a learned model’s predictive performance we used the Standardised Mean Squared Error (SMSE) on a held out test set of *N*_*\**_ individuals:

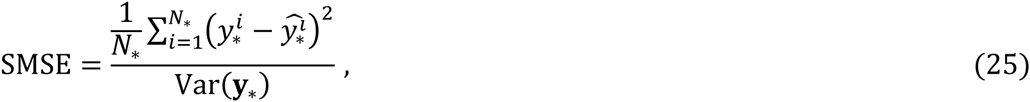
where 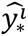 represents the predicted phenotype for the *i*^*th*^ individual and **y**_*\**_ is the vector of *N*_*\**_ test phenotypes. This is no more than the mean squared error normalised by the empirical variance of the data to enable more meaningful comparisons across datasets. The trivial method of predicting with the mean of the training data will have an SMSE of approximately 1.

### Judging significance - marginal Bayes factors

The marginal posterior distributions of the scale hyperparameters indicate relevance for the corresponding loci, but do not directly provide a measure of the significance of association^39^. Given the coding of alleles as 0 or 1, the form of the covariance function (3) indicates that, for a given model, only loci for which *τ*_*i*_∼𝒪(1)will contribute significantly. However, with limited data, there is likely to be considerable uncertainty over the underlying model. Therefore, we do not expect that the distributions of all of the *τ*_*i*_ corresponding to truly associated loci will concentrate sufficiently around such values for summaries such as the mean or median to satisfy this criterion. Therefore, we assessed the strength of evidence for the relevance of the *i*^*th*^ locus by computing a Bayes Factor (BF) for the hypothesis, *H*_1_: *τ*_*i*_ ≤ *ζ*, against the hypothesis, *H*_0_: *τ*_*i*_ > *ζ* for a given threshold, *ζ*.

The Bayes factor was obtained simply as the ratio of posterior and prior odds^25^:

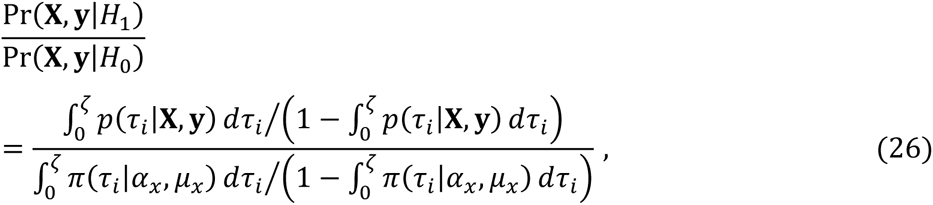
where *p*(.) and π(.) represent the posterior and prior densities respectively. Empirical estimates for 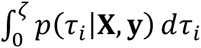 were obtained from the samples:

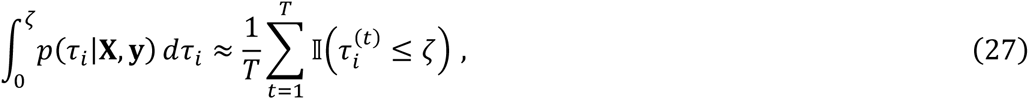
where the indicator function, 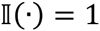 if its argument is true and 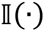 =0 otherwise.

In the case when a hierarchical prior was employed over *μ*_*x*_ we also estimated the implied prior from the samples:

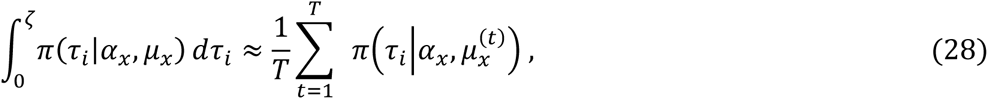

Values of 2, 6 and 10 for twice the natural logarithm of the Bayes Factor are commonly taken to indicate positive, strong and very strong evidence respectively^25^.

Although many choices of *ζ* are reasonable, for the analysis of the yeast data we chose 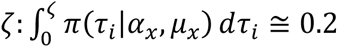 for each condition. For this choice, the threshold for positive evidence (2log(BF) < 2) was not reached by approximately 5% of the additive QTLs aggregated across all conditions. This was consistent with the reported, estimated False Discovery Rate for these additive QTLs8. In general, one can calibrate the threshold by repeating the GPR analysis on the same data but with the phenotype values permuted so that no true signals of association are expected. To be valid, however, the permutation must be done while accounting for the trait covariance structure. We used the MVNpermute R package to construct permuted datasets with this property. ^49^ Based on the output of such runs, a threshold can be chosen to ensure that (2log(BF) < 2) for all SNPs. We employed this approach for the analysis of the rat data. We found that choosing the threshold based on the scale below which 0.15 of the prior mass was located, 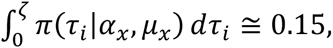 gave an appropriate threshold (**Supplementary Figure 23**). In this case, as we employed the hierarchical prior, samples of *μ*_*x*_ differed across runs. Therefore we used (28) to estimate the appropriate threshold for the actual and permuted data separately ensuring that the threshold, *ζ*, for the actual data met the same critierion 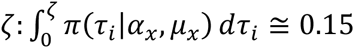 as that of the permuted data.

### Model Comparison

GPMM identifies SNPs that are important for explaining trait variance, possibly through interactions, but does not identify specific interaction effects. To examine the evidence for specific interactions between SNPS identified by GPMM as being associated with the rat phenotype, we computed maximum likelihood estimates (MLEs) for different nested models and performed standard likelihood ratio tests to generate p values (**Supplementary Table 4**). MLEs were computed using custom R code which implemented a previously described LMM algorithm^50^ which reduces estimation to a 1-dimensional optimization problem.

### 1-dimensional toy example

Samples were generated from a simple sine function with added Gaussian noise. For the *i*^*th*^ input, *x*_*i*_, we generated pairs of replicate samples, *y*_*ij*_ according to:

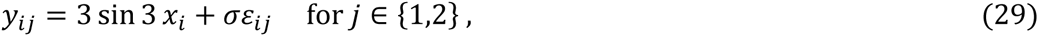
where 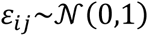 and *α* was chosen so that 80% of the variance was explained by the model. Samples were generated uniformly at random over a range 0 ≤ *x*_*i*_ ≤ 50 and subsets of size 30 were chosen as training data for learning.

## Acknowledgements

We are grateful to SURFsara (www.surfsara.nl) for use of the Lisa Compute Cluster. We are also grateful for use of the Darwin Supercomputer of the University of Cambridge High Performance Computing Service (http://www.hpc.cam.ac.uk/), provided by Dell Inc. using Strategic Research Infrastructure Funding from the Higher Education Funding Council for England and funding from the Science and Technology Facilities Council. KS moved from Nijmegen to Oxford during this work. The other authors are grateful to Prof. Jonathan Marchini for suggesting the analysis of the rat data, generously permitting KS time to perform the analysis and for supervising that analysis.

## Author Contributions

The analysis method was conceived and implemented by KS with contributions from WW. Analyses of the yeast data were designed by KS, WW, AA-V, BF, CA and HK. Analyses of the rat data were designed by JM and KS. Analyses were performed by KS. The manuscript was written by KS and incorporates comments from all the other authors.

## REFERENCES

1. Manolio, T.A. et al. Finding the missing heritability of complex diseases. Nature 461, 747–753 (2009).

2. Lander, E.S. Initial impact of the sequencing of the human genome. Nature 470, 187–197 (2011).

3. Visscher, P.M., Brown, M.A, McCarthy, M.I. & Yang, J. Five Years of GWAS Discovery. Am. J. Hum. Genet. 90, 7–24 (2012).

4. Moore, J.H. The Ubiquitous Nature of Epistasis in Determining Susceptibility to Common Human Diseases. Hum. Hered. 56, 73–82 (2003).

5. Phillips, P. C. Epistasis - the essential role of gene interactions in the structure and evolution of genetic systems. Nat. Rev. Genet. 9, 855–867 (2008).

6. Hemani, G., Knott, S. & Haley, C. An Evolutionary Perspective on Epistasis and the Missing Heritability. PLoS Genet. 9, e1003295 (2013).

7. Shao, H. et al. Genetic architecture of complex traits: Large phenotypic effects and pervasive epistasis. Proc. Natl. Acad. Sci. 105, 19910–19914 (2008).

8. Bloom, J. S., Ehrenreich, I. M., Loo, W. T., Lite, T. V. & Kruglyak, L. Finding the sources of missing heritability in a yeast cross. Nature 494, 234–237 (2013).

9. He, Xionglei, Qian, Wenfeng, Wang, Zhi, Li, Ying & Zhang, Jianzhi, Prevalent positive epistasis in Escherichia coli and Saccharomyces cerevisiae metabolic networks. Nat. Genet. 42, 272–276 (2010).

10. Chari S & Dworkin I The Conditional Nature of Genetic Interactions: The Consequences of Wild-Type Backgrounds on Mutational Interactions in a Genome-Wide Modifier Screen. PLoS Genet 9, (8): e1003661. (2013)

11. Culverhouse, R., Suarez, B.K., Lin, J. & Reich, T. A Perspective on Epistasis: Limits of Models Displaying No Main Effect. Am. J. Hum. Genet. 70, 461–471 (2002).

12. Zuk, O., Hechter, E., Sunyaev, S.R. & Lander, E.S. The mystery of missing heritability: Genetic interactions create phantom heritability. Proc. Natl. Acad. Sci. 109, 1193–1198 (2012).

13. Yang, J. et al. Common SNPs explain a large proportion of the heritability for human height. Nat. Genet. 42, 565–569 (2010).

14. Ritchie, M.D. et al. Multifactor-Dimensionality Reduction Reveals High-Order Interactions among Estrogen-Metabolism Genes in Sporadic Breast Cancer. Am. J. Hum. Genet. 69, 138–147 (2001).

15. Fang, G. et al. High-Order SNP Combinations Associated with Complex Diseases: Efficient Discovery, Statistical Power and Functional Interactions. PLoS ONE 7, e33531 (2012).

16. Nelson, M.R., Kardia, S.L.R., Ferrell, R.E. & Sing, C.F. A Combinatorial Partitioning Method to Identify Multilocus Genotypic Partitions That Predict Quantitative Trait Variation. Genome Res. 11, 458–470 (2001).

17. Zhang Y, Liu J: Bayesian inference of epistatic interactions in case-control studies. Nat. Genet., 39, 1167–1173 (2007).

18. Wu, J., Devlin, B., Ringquist, S., Trucco, M. & Roeder, K. Screen and clean: a tool for identifying interactions in genome-wide association studies. Genet. Epidemiol. 34, 275–285 (2010).

19. Kam-Thong, T., Pütz, B., Karbalai, N., Müller-Myhsok, B. & Borgwardt, K., Epistasis detection on quantitative phenotypes by exhaustive enumeration using GPUs. Bioinformatics 27, 214–221 (2011).

20. Lippert, C., et al., An Exhaustive Epistatic SNP Association Analysis on Expanded Wellcome Trust Data. Sci. Rep. 3, 1099 (2013).

21. Kirino, Y. et al., Genome-wide association analysis identifies new susceptibility loci for Behcet’s disease and epistasis between HLA-B[ast]51 and ERAP1. Nat. Genet., 45, 202–207 (2013).

22. Rasmussen, C.E. & Williams, C.K.I. Gaussian Processes for Machine Learning. MIT Press, Cambridge, Massachusetts, USA, (2006).

23. Duane, S., Kennedy, A.D., Pendleton, B.J., & Roweth, D. Hybrid Monte Carlo. Physics Letters B 195, 216–222 (1987)‥

24. Yang J., Lee S.H., Goddard M.E. & Visscher P.M. GCTA: a tool for Genome-wide Complex Trait Analysis. Am. J. Hum. Genet. 88, 76–82 (2011)

25. Kass, R. E. & Raftery, A.E. Bayes factors. J. Am. Stat. Assoc. 90, 773–795 (1995).

26. Baud, A. et al., Combined sequence-based and genetic mapping analysis of complex traits in outbred rats. Nat. Genet. 45, 767–775 (2013).

27. Zhou. X. & Stephens, M., Genome-wide efficient mixed-model analysis for association studies. Nat. Genet. 44, 821–824, (2012).

28. Bellou, A., & Finn, P. W. Costimulation: critical pathways in the immunologic regulation of asthma. Curr. Allergy Asthma Rep. 5, 149–154, (2005).

29. Choy, E. H. Selective modulation of T-cell co-stimulation: a novel mode of action for the treatment of rheumatoid arthritis. Clin. Exp. Rheumatol. 27, 510–518, (2009).

30. Shumilina, E. et al., Regulation of calcium signaling in dendritic cells by 1,25-dihydroxyvitamin D3. The FASEB Journal 24, 1989–1996 (2010).

31. Butte MJ, Lee SJ, Jesneck J, Keir ME, Haining WN, Sharpe AH. CD28 Costimulation Regulates Genome-Wide Effects on Alternative Splicing. PLoS ONE 7(6): e40032. (2012)

32. Yabas, M. et al. Differential Requirement for the CD45 Splicing Regulator hnRNPLL for Accumulation of NKT and Conventional T Cells. PLoS ONE 6(11): e26440. (2011).

33. Mackay, T., Epistasis and quantitative traits: using model organisms to study gene-gene interactions. Nat. Rev. Genet., 15, 22–33 (2014).

34. Mathew, B., et al. Bayesian adaptive Markov Chain Monte Carlo estimation of genetic parameters. Heredity, 109, 235–245 (2012).

35. Sawcer S., et al., A genome screen in multiple sclerosis reveals susceptibility loci on chromosome 6p21 and 17q22. Nat. Genet. 13, 464–468 (1996).

36. International Genetics of Ankylosing Spondylitis Consortium, Identification of multiple risk variants for ankylosing spondylitis through high-density genotyping of immune-related loci. Nat Genet. 45, 730–738 (2013).

37. Wang, A. et al., Gene–gene and gene–environment interactions in ulcerative colitis. J. Human Genetics 132, 547–558 (2013).

38. Dahl, A. et al. Multiple phenotype imputation for genetic studies. Nat. Genet. (2015) (in press)

## References

39. MacKay, D.J., Neural Networks and Machine Learning, Springer-Verlag, Berlin, Germany (1998).

40. Rasmussen, C.E. Evaluation of Gaussian Processes and other methods for non-linear regression. PhD Thesis, University of Toronto, Toronto, Canada (1996).

41. Zou, F., Huang, H., Lee, S. & Hoeschele, I. Nonparametric Bayesian Variable Selection With Applications to Multiple Quantitative Trait Loci Mapping With Epistasis and Gene–Environment Interaction. Genetics 186, 385–394 (2010).

42. Robert, C.P. & Casella, G. Monte Carlo Statistical Methods, Springer, New York, USA, (2004)

43. Horowitz, A.M., A generalized guided Monte Carlo algorithm. Physics Letters B 268, 247–252 (1991).

44. Girolami, M. & Calderhead, B. Riemann Manifold Langevin and Hamiltonian Monte Carlo Methods. JRSS B 73 123–214 (2011).

45. Gelman, A and Rubin, DB Inference from iterative simulation using multiple sequences, Statistical Science, 7, 457–511 (1992).

46. Speed, D., Hemani, G., Johnson, M.R and Balding, D.J. Improved Heritability Estimation from Genome-wide SNPs Am. J. Hum. Genet. 91, 1011–1021 (2012).

47. Bates, D., Maechler, M. & Bolker, B. lme4: Linear Mixed-Effects Models Using S4 Classes http://CRAN.R-project.org/package=lme4 (2011).

48. Endelman, J.B. Ridge regression and other kernels for genomic selection with R package rrBLUP. Plant Genome 4, 250–255 (2011).

49. Abney, M. Permutation testing in the presence of polygenic variation. Genet. Epidemiol. 39, 249–258 (2015).

50. Lippert, C. et al. FaST linear mixed models for genome-wide association studies. Nat. Meth. 8, 833–835 (2011).

